# CRISPR knockouts of *pmela* and *pmelb* engineered a golden tilapia by regulating relative pigment cell abundance

**DOI:** 10.1101/2021.12.09.471900

**Authors:** Chenxu Wang, Jia Xu, Thomas D. Kocher, Minghui Li, Deshou Wang

## Abstract

Premelanosome protein *(pmel)* is a key gene for melanogenesis in vertebrates. Mutations in this gene are responsible for white plumage in chicken, but its role in pigmentation of fish remains to be demonstrated. In this study we found that most fishes have two *pmel* genes arising from the teleost-specific whole genome duplication. Both *pmela and pmelb* were expressed at high levels in the eyes and skin of Nile tilapia. We mutated both genes in tilapia using CRISPR/Cas9 gene editing. Homozygous mutation of *pmela* resulted in yellowish body color with weak vertical bars and a hypo-pigmented retinal pigment epithelium (RPE) due to significantly reduced number and size of melanophores. In contrast, we observed an increased number and size of xanthophores in mutants compared to wild-type fish. Homozygous mutation of *pmelb* resulted in a similar, but milder phenotype than *pmela*^-/-^ mutants, without effects on RPE pigmentation. Double mutation of *pmela* and *pmelb* resulted in loss of additional melanophores compared to the *pmela*^-/-^ mutants, and also an increase in the number and size of xanthophores, producing a strong golden body color without bars in the trunk. The RPE pigmentation of *pmela*^-/-^*;pmelb*^-/-^ was similar to *pmela*^-/-^ mutants, with much less pigmentation than *pmelb*^-/-^ mutants and wild-type fish. Taken together, our results indicate that, while both *pmel* genes are important for the formation of body color in tilapia, *pmela* plays a more important role than *pmelb*. To our knowledge, this is the first report on mutation of *pmelb* or both *pmela;pmelb* in fish. Studies on these mutants suggest new strategies for breeding golden tilapia, and also provide a new model for studies of *pmel* function in vertebrates.

**Author Summary:** Melanophores, the most common pigment cell type, have been studied for nearly 150 years. Many genes are involved in melanoblast migration, melanophore differentiation, and melanin biosynthesis. *Pmel* is fundamental for melanosome development by directing melanin biosynthesis and melanosome phase transition. Specifically, PMEL can form a fibrillar structure within the melanosome upon which melanin is deposited. We identified two *pmel* genes in Nile tilapia arising from the teleost-specific whole genome duplication. Disruption of either *pmela* or *pmelb* in tilapia leads to significant hypo-pigmentation. PMEL disrupted fish showed not only a reduction in melanin and tiny melanophores, but also a significant increase in the number of xanthophores, and even guanine-filled melanophores, which led to a golden tilapia with hypo-pigmented RPE. Our study confirmed the role of *pmel* in melanin biosynthesis and maturation, and also highlighted its effects on melanophore number and size. These results provide new insights into pigment cell biology and will help us better understand the mechanisms of color patterning in teleosts. Knockout of both *pmela* and *pmelb* provide a new strategy for engineering a golden tilapia, which might provide a foundation for developing new strains in the tilapia industry.

## Introduction

Body color and pattern is of great significance for environmental adaptation and survival in vertebrates [1-3]. Some reports have also pointed out that teleost adaptation to extreme environments, such as cavefish and some deep-sea fish, is often associated with the degeneration of eyes and reduction in skin pigmentation. These changes are usually due to mutation or loss of pigment related genes [4-7]. For instance, the *mc1r* [8, 9] and *oca2* [4, 5, 10, 11], have been reported to be the major locus for pigment loss in some cavefish. In recent studies, *pmelb* has also been suggested to be closely linked with melanin loss in some cavefish and deep sea fish, as the amino acid sequence of this gene was significantly different among species [12]. These results indicate that *pmel* might be of great significance for teleost adaptation and evolution.

Melanophores are the most widely distributed and most important pigment cells in protecting animals from UV irradiation and oxidative stress [13, 14]. Many genes are important for melanin biosynthesis, including *pmel*, are regulated by the transcription factor *mitf* [15, 16, 17]. Studies in a variety of vertebrates indicate that *pmel* is fundamental for melanin maturation and deposition [18-21] and involved in many pigment cell diseases (hypo- or hyper-pigmentation skin diseases like albinism, vitiligo and melanoma). In studies of humans, *pmel* (also known as *silver*/*gp100*) is a key factor for vitiligo [22]. Mutations of this gene also result in abnormal pigmentation of the retinal pigment epithelium (RPE) and known as ocular pigment dispersion and pigmentary glaucoma [18]. Studies in mice indicate that variation in *pmel* affects the shape of melanosomes, with subtle effects on visible coat color [19, 23]. Variation in *pmel* is also associated with white skin, meat and even eggs in farmed chicken [20]. Most teleosts have two *pmel* genes arising from the teleost-specific genome duplication. In the cichlid fish *Haplochromis latifasciatus, pmela* shows significantly higher expression in black vertical bars than in light pigmented inter-bars, which suggests that *pmela* might be involved in bar formation [24]. Knockout of *pmela* in larvae zebrafish (*Danio rerio*) causes defects in eye development and pigmentation of the RPE and the skin [18]. To date, no loss of function studies have been conducted in teleosts of *pmelb* single and *pmela;pmelb* double mutation.

There are four phases in melanosome development. First, phase I melanosomes develop from endosomal membranes. The formation of PMEL fibrils gradually converts phase I to phase II melanosomes. Only after the fibrillar structures are formed does melanin begin to be deposited to create phase III melanosomes. The mature phase IV melanosome has a high concentration of melanin. Melanin biosynthesis is directly controlled by genetic factors whose expression is regulated by the melanocyte-inducing transcription factor *mitf*, including the tyrosinase family genes, the iron channel *oca* family genes, the *bloc* family genes, the *slc* family genes, the *hps* family genes and the *pmel* genes. In addition, cell intrinsic environmental factors like the cell membrane electrification, pH levels and the concentration of metal ions (Na^+^, K^+^, Ca^2+^, Mg^2+^, etc.), also have critical effects on melanin synthesis [13, 14]. PMEL has been acknowledged to play the similar role with tyrosinase and HPS family members, by affecting the transition of melanosomes from phase I to II [21]. However, even though some studies have confirmed the role of *pmel* in tetrapod body color formation, mutations in this gene have been found to not always have the same effects on melanin biosynthesis and melanosome shape in different vertebrates [19, 20].

There are three major types of pigment cells responsible for teleost color patterning: the black/grey melanophores, the yellow/reddish xanthophores and the blue/purple/white iridophores. In zebrafish these three types of pigment cells are fundamental for stripe formation, through run and chase cell-cell interactions [25]. Recent studies on *Danio* pigment cells suggested that two groups of white pigment cells are critical for establishing pattern. White leucophores arises by trans-differentiation of adult melanophores. A second white cell type develops from a yellow-orange xanthophores or a xanthophore-like progenitor [26]. Similar studies in cichlids also demonstrated the importance of cell interactions in pigment patterning, even though the specific mechanisms might be different [24, 27-31]. Additionally, studies on rainbow trout suggested that homozygous mutation of an albinism gene, led to not only reduced number of melanophores, but also significantly larger size of xanthophores, which finally led to yellow-albino fish with golden color [32]. However, it remains to be further proved whether the change of the number, morphology and pigment synthesis of one type of pigment cells will also cause the corresponding change of the number, morphology and pigment synthesis of other pigment cells.

Disruption of PMEL in chicken led to complete white chicken, which was favored by consumers all over the world [20]. In Japanese quail, this gene was also associated with the formation of yellowish plumage through GWAS mapping [21]. In fish, the role of *pmel* in body color formation remains to be illustrated. Even though *pmela* has been acknowledged to be fundamental for melanin biosynthesis in larval zebrafish [18], detailed role of *pmela* and *pmelb* in teleost body color formation remains to be investigated. As a food fish farmed worldwide, tilapia has been favored for its strong resistance to disease, fast growth and high protein content. Tilapia is also an excellent model for scientific research because it has a published genome, short time to sexual maturity, short spawning cycle and large brood sizes. In a previous study, we developed the Nile tilapia as a model for studying teleosts color patterns, and identified severe vitiligo-like phenotypes in *pmel* F0 mutants, suggesting that *pmel* genes are indeed fundamental for tilapia body color formation [33]. However, whether the homozygous mutation of *pmel* genes will give a healthy attractive white, silver, yellowish, golden body color or even a confusing mosaic pattern in tilapia still remains to be investigated.

In this study, we characterized the evolutionary history of the *pmela* and *pmelb* genes, and analyzed their expression in Nile tilapia. We then used CRISPR/Cas9 gene editing to disrupt *pmela* and *pmelb* and create homozygous mutants. We also analyzed the phenotype of these mutants to infer the function of *pmel* in this species. Disruption of PMEL yielded a complete golden body color in Nile tilapia, which was reflected by the changes of pigment cell numbers and cell sizes of melanophores and xanthophores. Our study provides a path to engineer a new strain of golden tilapia and might also deepen the understanding of pigment cell biology and color pattern formation in teleosts.

## Results

### *Pmela* and *pmelb* diverged after the teleost-specific genome duplication

As shown in Figure 1, we found that euteleosts have two copies of *pmel* due to the third round of genome duplication, while non-bony fish and tetrapods have only one copy. Nile tilapia *pmela* was most similar to the zebra cichlid, followed by Japanese medaka, guppy and torafugu. The *pmelb* was also most similar to the zebra cichlid, followed by torafugu/tongue sole, guppy/Japanese medaka, and then other teleosts. The promoters of *pmela* and *pmelb* each contain several binding sites for *mitf*, consistent with the idea that both genes are directly regulated by *mitf* (Fig. S1 and S2). The closest ortholog to PMEL is Glycoprotein Nonmetastatic Melanoma Protein B (GPNMB), another transmembrane glycoprotein also regulated by *mitf* [34, 35].

**Fig 1.**
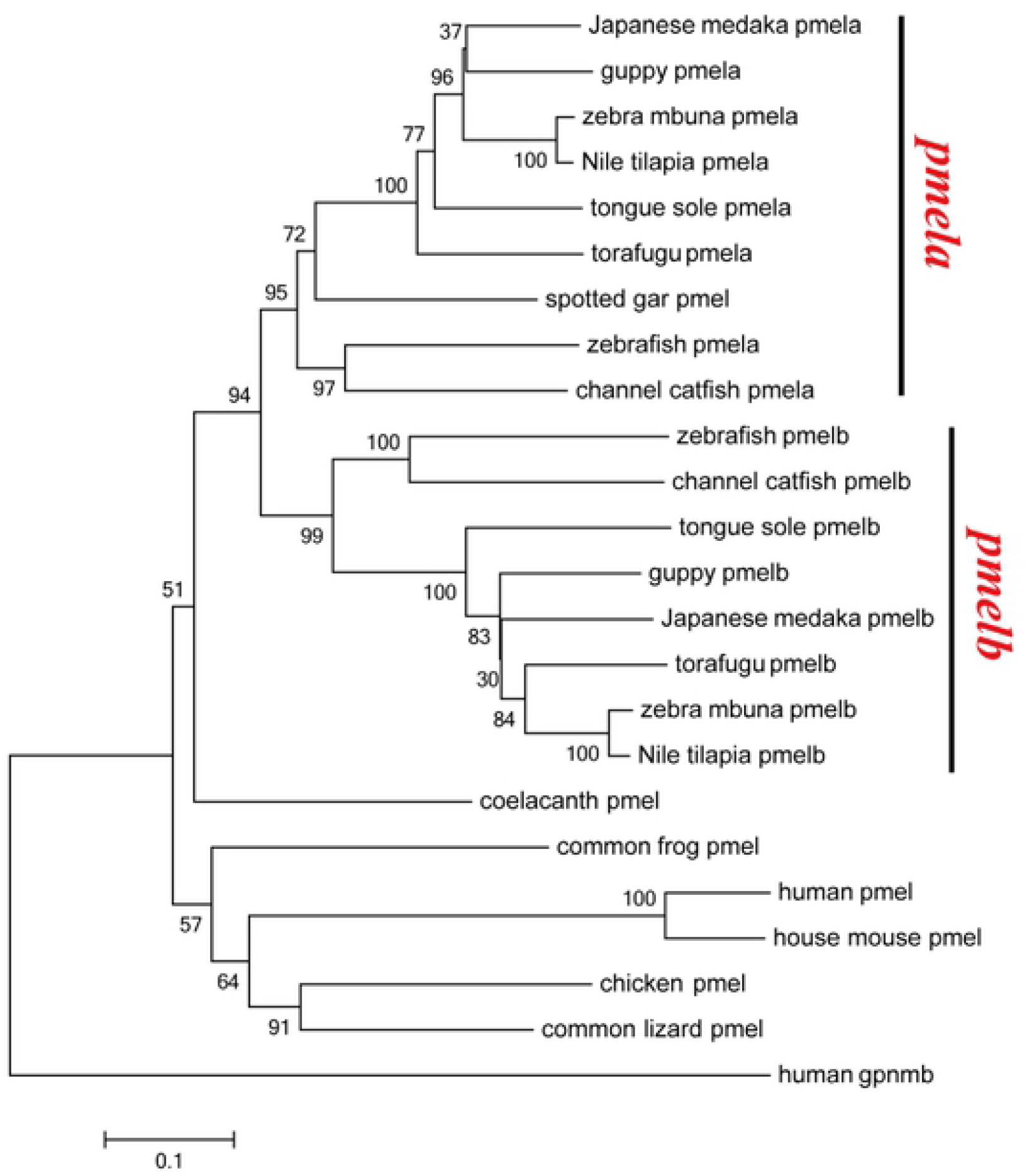
Phylogenetic tree of pmel genes from 15 representative species. The NJ method was used to construct the tree by MEGA6.0 software, using human GPNMB protein (the homologue of PMEL) as the out-group.. The multiple alignments of software Bioedit was used to align the amino acid sequences. The pmel genes were divided into 2 branches: pmela and pmelb. Eubony fish have two copies due to the third round of genome replication, while non bony fish and quadrupeds do not have two copies. The pmel amino acids were downloaded from zebra mbuna (XP_004545057 of pmela and XP_004560330 of pmelb, respectively), Nile tilapia (XP_019214528 of pmela and XP_003457123 of pmelb, respectively), Japanese medaka (XP_004069166 of pmela and XP_004071062 of pmelb, respectively), guppy(XP_008408226 of pmela and XP_008412883 of pmelb, respectively), torafugu (XP_029683274 of pmela and XP_029690408 of pmelb, respectively), channel cat fish (XP_017306583 of pmela and XP_017342597 of pmelb, respectively), spotted gar (XP_015199523), zebrafish (XP_021335225 of pmela and XP_691276 of pmelb, respectively), tongue sole (XP_024915820 of pmela and XP_008317243 of pmelb, respectively), coelacanth (XP_005986276), common frog (XP_040197009), human (NP_001186983), house mouse (XP_030100827), chicken (XP_040510678) and common lizard (XP_034959828).

### Expression of *pmela* and *pmelb* in Nile tilapia

RT-PCR analysis showed high levels of expression of both *pmela* and *pmelb* in eyes, skin and whole embryos. Weak expression of *pmela* was detected in heart and testis, but it was rarely expressed in other tissues (Fig S3). Some expression of *pmelb* was detected also in brain and fins (Fig S4). Across the different developmental stages (2, 4, 6, 8, 10, 20 and 30 dpf of the whole fish, 60, 90 dpf and adult of the black skin) of wild-type Nile tilapia, *pmela* and *pmelb* were detected with similar expression levels and trends at all stages. They were highly expressed at 4, 6, 8, 10 and 90 dpf, but showed less expression at 20, 30, 60 dpf and the adult stage. Neither gene was expressed at the 2 dpf blastula stage, probably because pigmentation genes are only expressed after neural crest cells (NCCs) are specified. Additionally, *pmela* and *pmelb* were consistently expressed in whole fish from 4-10 dpf, with the highest expression at 8 and 10 dpf (Fig S5). This was inconsistent with the observation of several waves of melanophore increase in our previous study, including key events at 5 dpf (the time melanophores spread on the yolk sac), 7 dpf (the time melanophores spread on both the trunk and the yolk sac), 12 dpf (the time melanophores spread across the whole trunk but are not detected in the fins) and 90 dpf (the time vertical bar patterns began to form in both trunk and fins) [33].

### Establishment of the *pmela*^-/-^, *pmelb*^-/-^ and *pmela*^-/-^*;pmelb*^-/-^ mutants in tilapia

Two gRNAs, targeting exon 4 of *pmela* and exon 3 of *pmelb*, were used to disrupt *pmela* and *pmelb* simultaneously in tilapia. The sequences of gRNA targets were CCCACCAAACCAGACGGTGCTCC and GGAAAAGTGACGTTTAATGTTGG, in which CCC and TGG were used as the PAM regions (Fig 2A-2C and 2G-2I). *Fsp*EI and *Tsp*45I respectively were used for enzymatic digestion of the amplified target regions.

**Fig 2.**
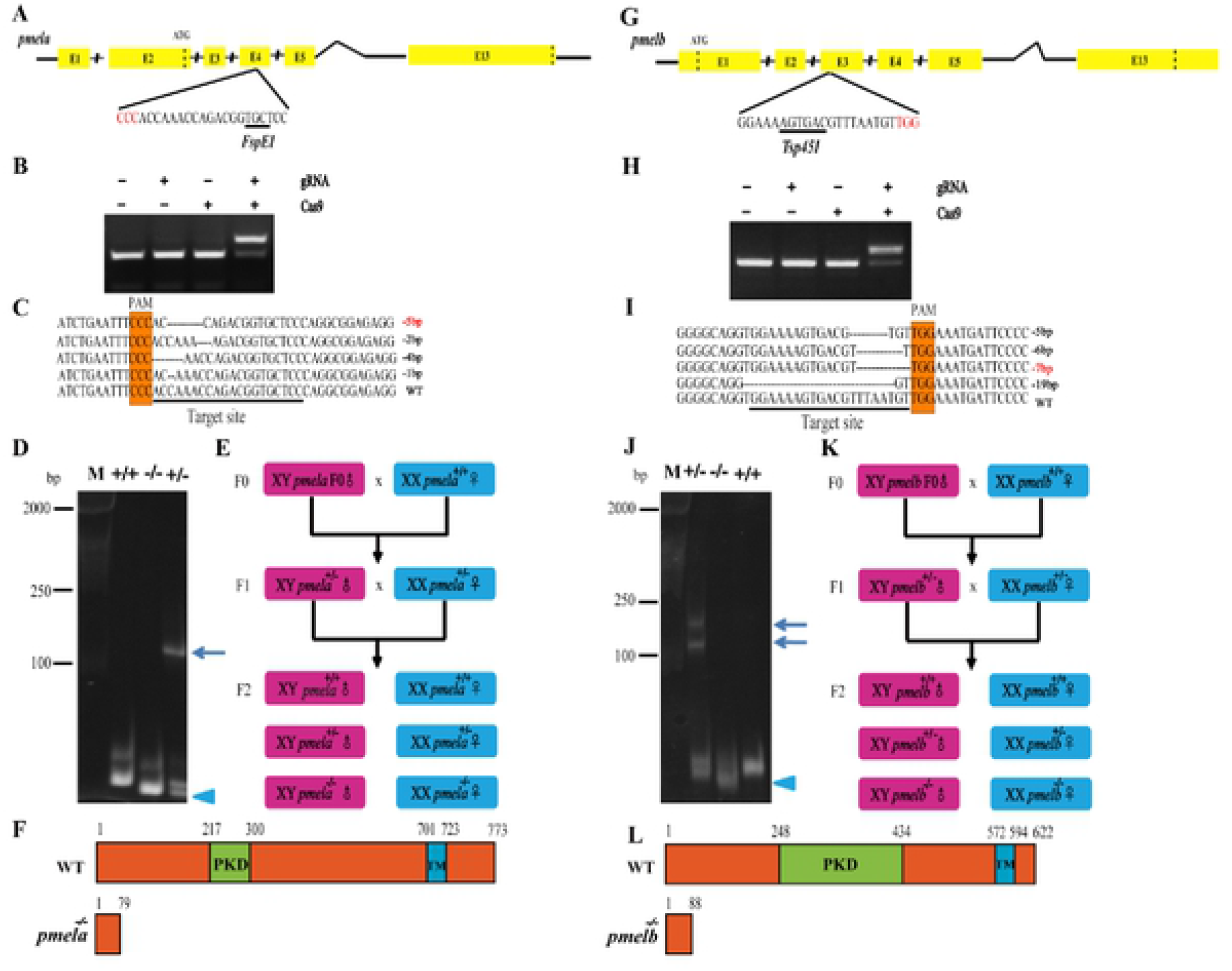
Establishment of pmela^-/-^ (A-F) and pmelb^-/-^ (G-L) mutant lines. A, G: Gene structures of pmela and pmelb showing the target site and the FspEI and Tsp451 restriction site, respectively. B, H: Restriction enzyme digestion of the amplified fragment of pmela and pmelb using primers spanning the target sites. The Cas9 mRNA and gRNA were added as indicated. C, I: Sanger sequencing results from the uncleaved bands were listed. The PAM is marked in orange. Deletions are marked by dashes (-) and numbers to the right of the sequences indicate the loss of bases for each allele. WT, wild type. D, J: Identification of pmela and pmelb F2 genotypes by hetero-duplex motility assay. Arrowheads show homo-duplexes and arrows show hetero-duplexes. E, K: Schematic diagram showing the breeding plans of pmela and pmelb F0 to F2 fish. F, L: Tilapia pmela and pemelb showing the functional domains (orange) including PKD domain (green) and TM domain (blue).

F0 founders were screened by restriction enzyme digestion and Sanger sequencing (Fig 2B, 2C, 2H and 2I). The *pmela* and *pmelb* mutant fish with a high mutation rate (over 75%) were raised to sexual maturity and mated with wild-type tilapia to create F1 fish. In the *pmela* and *pmelb* F0 chimeras, various levels of hypo-pigmentation were observed in body and the iris was also hypo-pigmented, as we previously reported [33]. These results indicated that the two genes are fundamental for body color formation and eye pigmentation in tilapia, and are probably necessary for melanin synthesis in all melanophores (NCCs-derived and optic-cup derived melanophores).

The F1 mutant fish were obtained by crossing an F0 XY male with a wild-type XX female. Heterozygous *pmela* F1 offspring with a -5 bp deletion in the fourth exon were selected to breed the F2 generation (Fig 2C-2E). Likewise, heterozygous *pmelb* F1 offspring with a -7 bp deletion in the third exon were selected to breed an F2 generation (Fig 2I-2K). Finally, *pmela;pmelb* F2 were produced by mating a male *pmela* -5 bp F1 heterozygous mutant with a female -7 bp F1 *pmelb*-positive fish, then heterozygous *pmela;pmelb* F2 offspring with a -5 bp deletion in *pmela* and -7 bp deletion in *pmelb* were selected to breed the F3 generation (Fig S6). A heteroduplex mobility assay identified the heterozygous *pmela*^*+/-*^, *pmelb*^*+/-*^ and *pmela*^*+/-*^*;pmelb*^*+/-*^ individuals as those possessing both heteroduplex and homo-duplex amplicons vs. the *pmela*^+/+^ and *pmela*^-/-^ individuals with only homo-duplex amplicons (Fig 2D and 2J).

### Melanophore size, number and melanin biosynthesis were decreased in *pmel* mutants

Several waves of increase in melanophore number were detected in wild-type tilapia, during which melanophore populations increased to pattern the whole fish [33]. Like the tetrapods, melanophores in teleosts were also able to release melanosomes extracellularly (Fig S7). We chose to analyze the melanophores of wild-type and *pmel* mutant fish at 7 dpf, shortly after melanophores first appear on the embryo and spread on both the yolk sac and the trunk. The wild-type fish had many well-pigmented macro-melanophores and normal sized melanophores on the head from 7 dpf (Fig 3A and 3A’). However, pigmented melanophores were greatly reduced in the *pmela*^-/-^ (Fig 3B’ and 3B’) and *pmela*^-/-^*;pmelb*^-/-^ mutants (Fig 3D and 3D’). The remaining melanophores were unable to synthesis mature melanosomes, and most of the melanosomes were in phase II. The remaining macro-melanophores were light grey, a hypo-pigmented phenotype (Fig 3A, 3A’, 3D and 3D’). In contrast, although the *pmelb*^-/-^ mutants also had fewer melanophores, some melanophores were still able to develop dark, mature melanosomes (Fig 3B and 3B’). The number of melanophores in *pmela*^-/-^;*pmelb*^/-^ and *pmela*^-/-^*;pmelb*^-/-^ mutants was significantly lower than that of the wild-type fish. Although *pmelb*^-/-^ mutants retained more melanophores, most of the pigmented melanophores were fragmented or found in a state of pigment-aggregation. The *pmela*^-/-^*;pmelb*^-/-^ double mutants had significantly reduced melanophore numbers and sizes compared with both wild-type fish and the single gene mutants (Fig 3E and 3F). Additionally, xanthophores were detected in significantly higher numbers and larger sizes than the wild-type fish at 12 dpf, which gave the fish a yellowish head (Fig 4A-4C). This phenomenon probably could explain the complete golden color of the whole fish at later stages.

**Fig 3.**
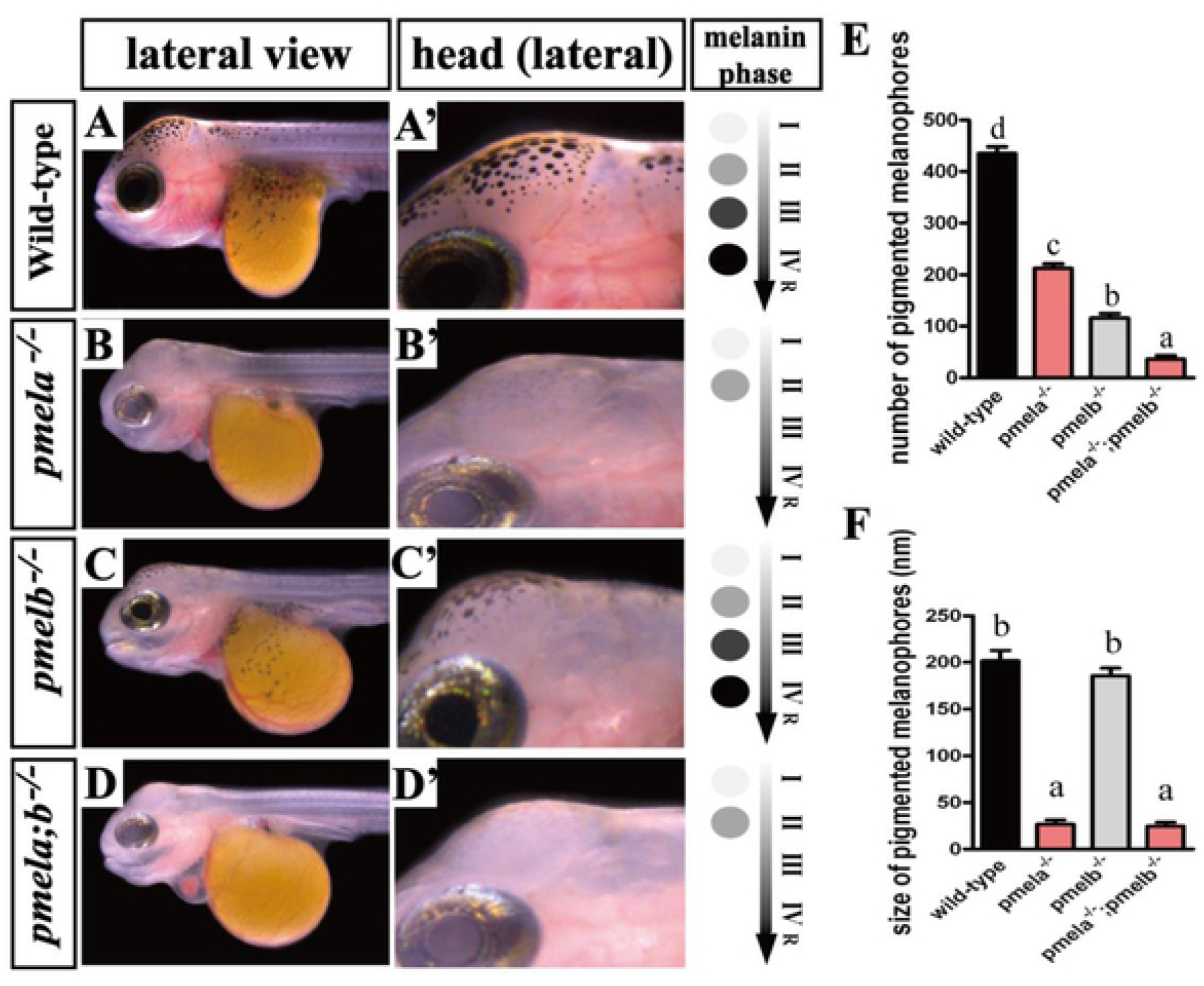
Mutation of pmela and pmelb resulted in significant melanophores reduction, cell size decrease and melanin biosynthesis degeneration at 7 dpf. In pmela^-/-^ and pmela^-/-^;pmelb^-/-^ mutants, the melanophores were heavily reduced, but the remaining melanophores (mainly macro-melanophores) still kept the stretch like outlooks, and everyone of them displayed hypo-pigmentation. Most of the melanin was in stage II. In wild-type fish and pmelb^-/-^ mutants, the melanophores were capable of biosynthesizing mature melanin until stage IV, and then release to further patterning the whole fish. And the melanophores in pmelb^-/-^ mutants were also heavily reduced when compared with the wild-type fish. The pigmented melanophores on the kept decrease from pmela^-/-^ -pmelb^-/-^-pmela^-/-^;pmelb^-/-^ mutants, and the difference was significant (P < 0.05). However, when related to the size of melanophores, there was no such an orderly decrease. The size of pmela^-/-^ and pmela^-/-^;pmelb^-/-^ mutants were significantly smaller than that of pmelb^-/-^ and wild-type fish. The single melanophore was unable to synthesize much melanin due to the size reduction and PMEL disruption. Data are expressed as mean ± SD (n=9). Significant differences in the data between groups were tested by one-way ANOVA and Duncan’s post-hoc test. P<0.05 was considered to be statistically significant, as indicated by different letters above the error bar. R, release.

**Fig 4.**
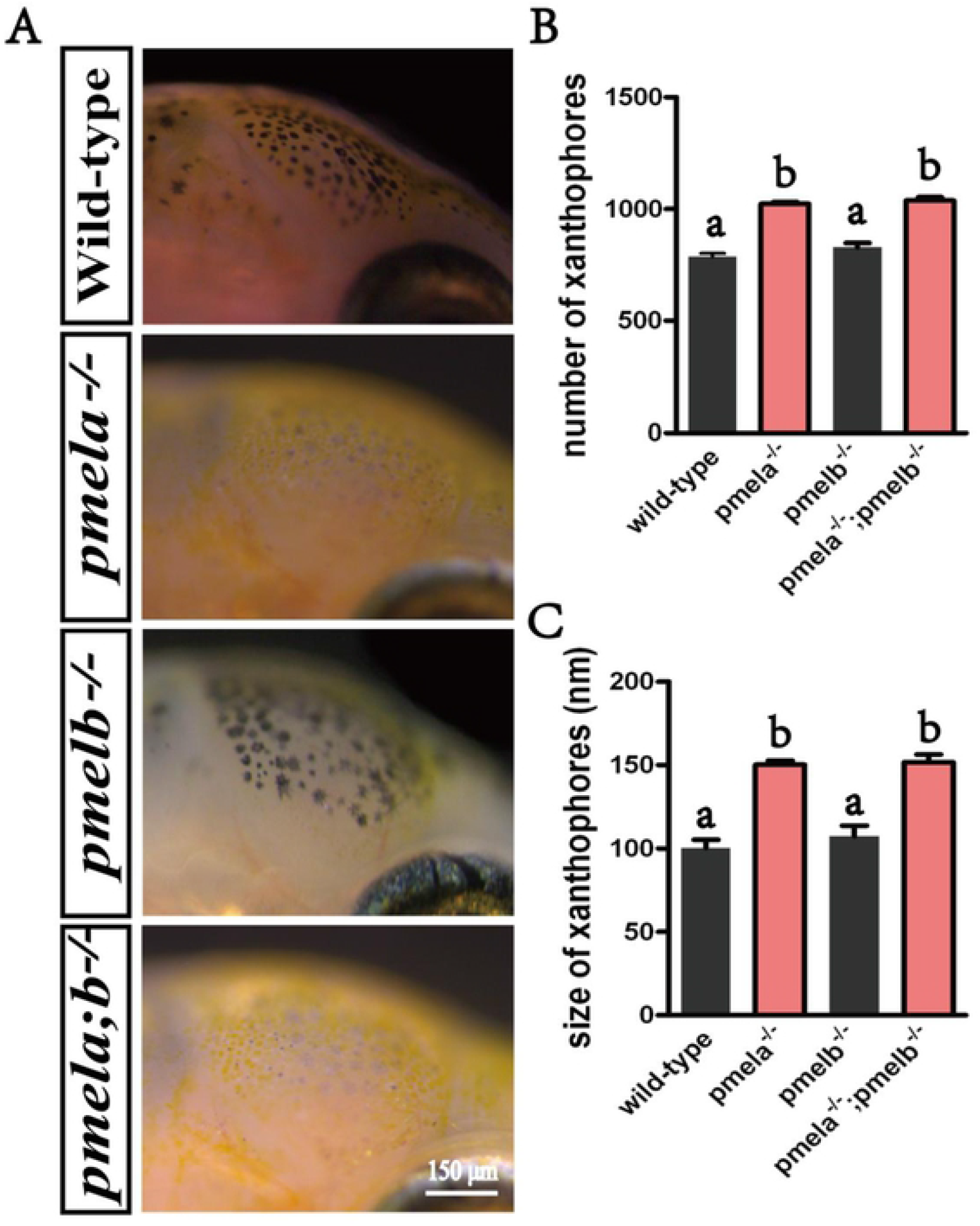
Mutation of pmela and pmelb resulted in melanophores reduction, melanin biosynthesis degeneration, more and larger sized xanthophores on the head at 12 dpf. A: Mature melanin inside the functional stretch-like melanophores was easily detected in the wild-type fish at 12dpf. However, the light grey melanophores with serious hypo-pigmentation were detected in the head of pmela^-/-^ and pmela^-/-^;pmelb^-/-^ mutants, the melanin inside were almost all in phase I or II so far. Thus melanophores were unable to biosynthesis mature melanin to further pattern the whole fish. Additionally, melanophores were heavily reduced. As a contrast, more and larger sized xanthophores were detected, which made the head a golden outlook. B and C: The number of xanthophores in pmela^-/-^ and pmela^-/-^;pmelb^-/-^ mutants were significantly higher than that of pmelb^-/-^ and wild-type fish, and the size of xanthophores in pmela^-/-^ and pmela^-/-^;pmelb^-/-^ mutants were significantly lager than that of pmelb^-/-^ and wild-type fish. Data are expressed as mean ± SD (n=5). Significant differences in the data between groups were tested by one-way ANOVA and Duncan’s post-hoc test. P<0.05 was considered to be statistically significant, as indicated by different letters above the error bar.

### Xanthophore number and size were increased in *pmel* mutants

In our previous study, we found that in wild-type fish xanthophores arose at 6 dpf, and sharply increased in number by 12 dpf [33]. We used 12 dpf wild-type, *pmela*^-/-^, *pmelb*^-/-^ and *pmela*^-/-^*;pmelb*^-/-^ mutants to analysis xanthophore number and size in PMEL disrupted fish. The *pmela*^-/-^ and *pmela*^-/-^*;pmelb*^-/-^ mutants showed significantly more and larger sized xanthophores, which was a major reason for the yellowish color of those mutants (Fig 4A-4C). In wild-type fish, the xanthophores were often in a pigment-aggregated state, with a much smaller size than the expanded melanophores [33]. However, in the *pmela*^-/-^ and *pmela*^-/-^ *;pmelb*^-/-^ mutants, the lateral top of the heads showed lots of yellow pigmentation, the xanthophores were filled with pteridines/carotenoids, and the sizes were much larger than wild-type fish and *pmelb*^-/-^ mutants. The remaining melanophores were small and hypo-pigmented with small sizes, similar to the 7 dpf mutants (Fig 4A). These results indicated that loss of pigment biosynthesis function in melanophores led to decreased melanophore number and size, and a corresponding increase in xanthophore size and number.

### RPE, iris pigmentation and eye development were affected in *pmela* mutants, while only iris pigmentation was affected in *pmelb* mutants

Reduced eye pigmentation was observed in all developmental stages of *pmel* mutants, and the *pmela*^-/-^ and *pmela*^-/-^*;pmelb*^-/-^ mutants were free of melanin in the eyes at early larvae stages (before 60 dpf). Thus fish at 60 dpf and 90 dpf were used as materials for studying eye pigmentation and development. Pigmentation in the eyes (including both iris and RPE) displayed significant differences between the mutants and wild-type fish. The *pmela*^-/-^ and *pmela*^-/-^*;pmelb*^-/-^ mutants showed significant hypo-pigmentation in eyes, the RPE was dark red, and the iris was golden (Fig 5B and 5D). In contrast, the wild-type fish and *pmelb*^-/-^ mutants had a black RPE, and a normally pigmented iris (Fig 5A and 5C). The development of eyes was also altered in *pmela*^-/-^ and *pmela*^-/-^*;pmelb*^-/-^ mutants. Around 1/3 of both *pmela*^-/-^ and *pmela*^-/-^*;pmelb*^-/-^ mutants had both serious reduction of iris iridescent-white pigmentation and significant eye abnormal development. Pigment deposition onto the inner surface of the cornea causes the appearance of the Krukenberg spindle. The area of iris covered by melanin stuck to the cornea was around 1/3 compared with the central black RPE area (Fig 5B’, 5D’ and 5E). Even though slow restoration of RPE pigmentation was detected at 90 dpf, part of the RPE area still had a blood red color indicative of insufficient melanin, and the Krukenberg spindle phenotype was durable (Fig 5E). In contrast, the wild-type fish and *pmelb*^-/-^ mutants showed no significant abnormalities of eye development (e.g. pigment stuck to the cornea), even though the *pmelb*^-/-^ mutants hypopigmentation of the eyes. The results of global pigmentation in eyes were in consistent with the phenotype we showed above (Fig 5F).

**Fig 5.**
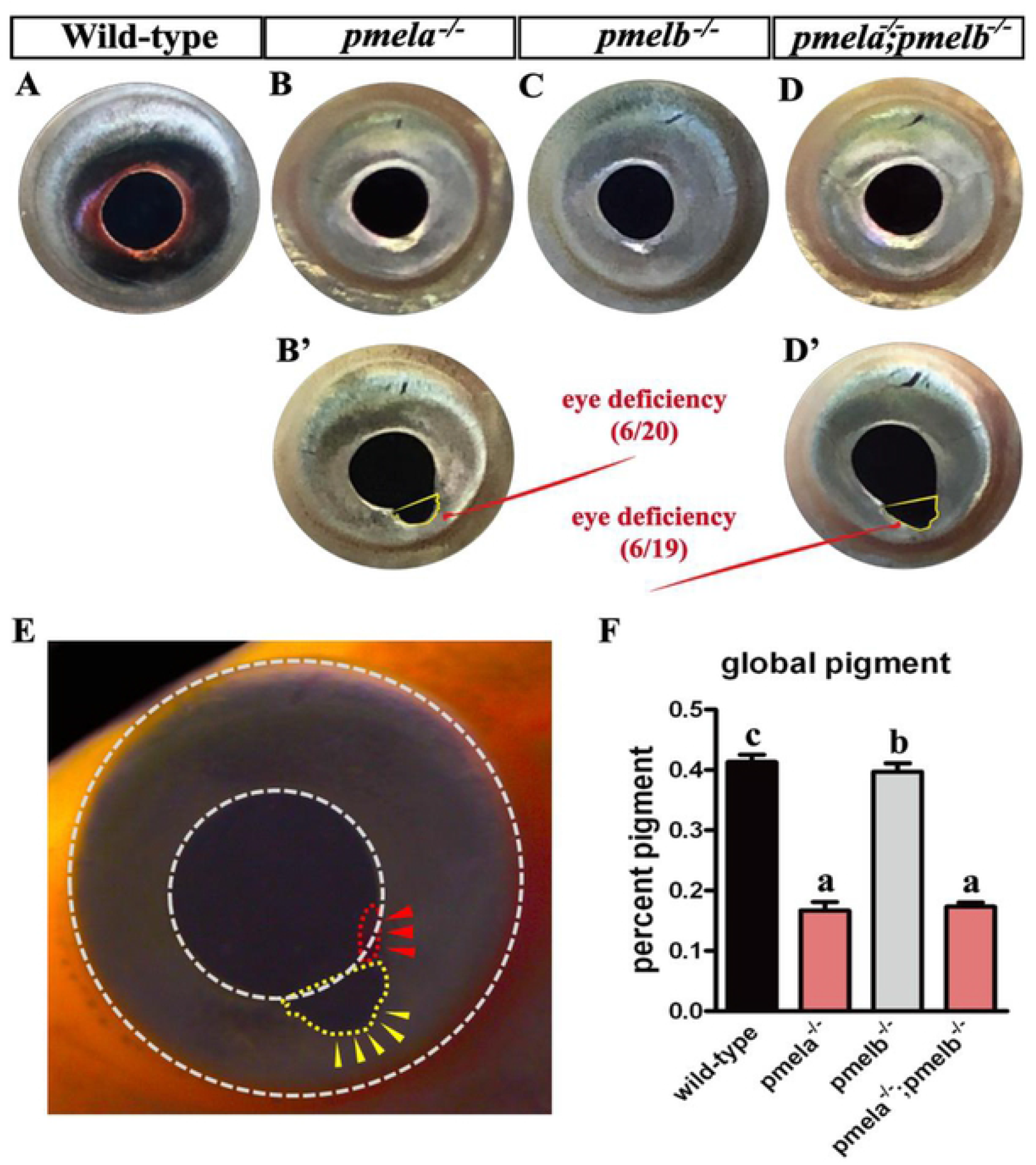
Mutation of pmela resulted in RPE hypo-pigmentation and eye deficiency at 60 and 90 dpf. A-D: The pmela^-/-^ and pmela^-/-^;pmelb^-/-^ mutants showed deep red color in RPE, but difficult to be distinguished from the lateral view. While deep black color was detected in the RPE of wild-type fish and pmelb^-/-^ mutants. No black pigmentation was detected in the iris of pmela^-/-^ and pmela^-/-^;pmelb^-/-^ mutants, but some black pigmentation was detected in the iris of pmelb^-/-^ mutants. B’ and D’: Additionally, the pmela^-/-^ (6/20) and pmela^-/-^; pmelb^-/-^ (6/19) mutants were detected with serious eye deficiency (ocular pigment dispersion and pigmentary glaucoma) in aroud 1/3 mutants, compared with wild-type fish and pmelb^-/-^ mutants at 60 dpf. E: Around 1/3 of the pmela^-/-^ and pmela^-/-^;pmelb^-/-^ mutants were detected with abnormal eye development at 90 dpf. Reflected by some melanin spread out of the round iris (yellow dashed box highlighted by yellow arrow heads). Additionally, even though slow restoration of melanin biosynthesis was observed with individual development, a part of the RPE and iris area was detected with almost no melanin, reflected by blood red color (red dashed box highlighted by red arrow heads, under strong bright field). Generally, this is a phenotype of Krukenberg spindle (pigment stuck to the cornea). F: The statistical analysis of eye pigmentation, the percentages of pmela^-/-^ and pmela^-/-^;pmelb^-/-^ mutants were significantly lower than in pmelb^-/-^ mutants and wild-type fish (p>0.05). And there was also significant difference between pmelb^-/-^ mutants and wild-type fish (p<0.05). Data are expressed as mean ± SD (n=9). Significant differences in the data between groups were tested by one-way ANOVA and Duncan’s post-hoc test. P<0.05 was considered to be statistically significant, indicated by different letters above the error bar.

### Melanophore number and melanin synthesis were restored in older *pmela*^-/-^ and *pmela*^-/-^ *;pmelb*^-/-^ mutants

Phenotypic analysis of the caudal fins suggested that pigmented melanophores increased with age in the mutants. The *pmela*^-/-^ and *pmela*^-/-^*;pmelb*^-/-^ mutants had only a few pigmented melanophores before 60 dpf. However, after further development, small pigmented melanophores were observed in the *pmel* mutants. The caudal fins in wild-type, *pmela*^-/-^, *pmelb*^-/-^ and *pmela*^-/-^*;pmelb*^-/-^ mutants were used as samples to study the re-pigmentation and re-patterning progress of melanophores. In wild-type fish at 60 dpf, hyper-pigmented bars separated by light colored inter-bars were observed in the caudal fin (Fig 6A). Melanophores were detected in large numbers in the hyper-pigmented bar regions (Fig 6A’). In *pmela*^-/-^ mutants at 60 dpf, yellowish caudal fins without any bars were detected (Fig 6B). No pigmented melanophores were detected at this time period. What was striking was that aggregated iridescent purple guanine was detected in cells with branching-like-clusters sharing a shape and size similar to melanophore, both in caudal fins and even scales (Fig 6B’ and Fig S8). We suggest that melanophores under specific conditions (melanin-free or melanin biosynthesis deficient) were able to synthesize or accumulate guanines. In *pmelb*^-/-^ mutants at 60 dpf, no obvious bars but some pigmented melanophores were detected in the caudal fins (Fig 6C and 6C’). In *pmela*^-/-^*;pmelb*^-/-^ mutants at 60 dpf, the whole fins were yellowish without bars, and many purple-guanine-pigmented melanophores were detected randomly spread across the whole fins (Fig 6D and D’). At 90 dpf, a significant increase in pigmented melanophores was detected in the caudal fin (Fig 6E). Additionally, many red erythrophores/xanthophores were detected in the fins (Fig 6E’). In *pmela*^-/-^ mutants at 90 dpf, a significant increase in small melanophores was detected across the whole fins, and they spread evenly across the fin (Fig 6F). Large numbers of evenly distributed white iridophores were also detected (Fig 6F’). In *pmelb*^-/-^ mutants at 90 dpf, increased numbers of melanophores were detected in the caudal fins, with larger size and significant higher number than the *pmela*^-/-^ mutants at the same developmental stage (Fig 6G). Additionally, the white iridophores also increased compared with the wild-type fish, similar to the situation revealed in *pmela*^-/-^ mutants. In *pmela*^-/-^*;pmelb*^-/-^ double mutants at 90 dpf, melanophores were also more abundant compared to the 60 dpf *pmela*^-/-^*;pmelb*^-/-^ mutants. The melanophores in the double mutants were in spot-like small size, and fewer in number than in wild-type, *pmela*^-/-^ and *pmelb*^-/-^ mutants at the same stage. Interestingly, the number of red erythrophores/xanthophores increased in the double mutants, which contributes to a more yellowish or reddish body color (Fig 6H, H’), and the whole fish were seriously hypo-pigmented compared to the wild-type fish (Fig S9). Statistical analysis of the melanophores of the *pmel* mutants and wild-type fish at 60 and 90 dpf was consistent with the descriptions above (Fig 6I and 6J). The restoration of melanin biosynthesis in *pmela*^-/-^ and *pmela*^-/-^*;pmelb*^-/-^ mutants at older ages indicated that additional pathways might be involved in melanin biosynthesis.

**Fig 6.**
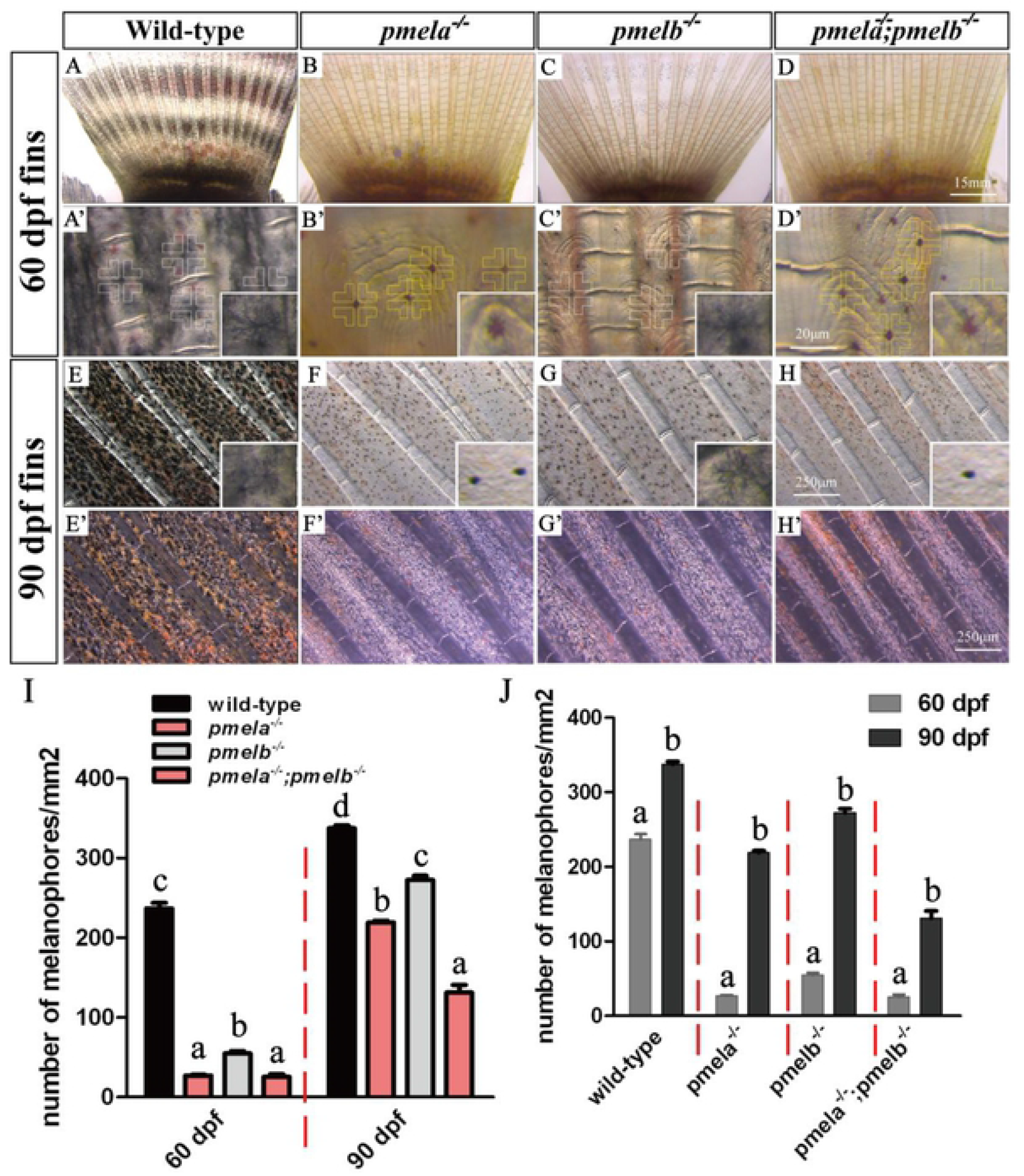
Mutation of pmela and pmelb resulted in melanophores reduction at 60 dpf, but with some restoration at 90 dpf in caudal fins. A, A’, E and E’: Large number of melanophores, erythrophores and xanthophores were observed in 60 and 90 dpf wild-type caudal fin. Obvious black vertical bars and light pigmented inter-bars were also detected in wild-type fish, the melanophores with branch like clusters (white cross) were in a higher density in bars than in inter-bars. B, B’, F and F’: Almost no pigmented melanophores were detected in the caudal fins of pmela^-/-^ mutants at 60 dpf, but xanthophores were detected. Some guanine filled in the not fully pigmented melanophores, thus making it showing branching-like clusters with red and purple colors (golden cross). The pigmented melanophores were detected with spot-like shape in 90 dpf pmela^-/-^ mutants. The iridophores were most in white color, and many xanthophores were also detected. Both the latter two types of pigment cells were widely distributed between the fin rays. C, C’, G and G’: A few pigmented melanophores were detected in the fins of pmelb^-/-^ mutants at 60 dpf, with branching-like clusters. Many iridophores and xanthophores were also detected. More pigmented melanophores were detected in pmelb^-/-^ mutants at 90 dpf than at 60 dpf. All these melanophores were in a smaller size than the wild-type melanophores, but larger than in pmela^-/-^ mutants. Additionally, xanthophores and iridophores were detected with the similar color patterning with pmela^-/-^ mutants at the same developmental stage. D, D’, H and H’: Completely no pigmented melanophores were detected in the caudal fins of pmela^-/-^;pmelb^-/-^ mutants. While many iridophores were observed to be filled in the not pigmented melanophores, thus showing red and purple colors (golden cross). Significant less melanophores than wild-type, pmela^-/-^ and pmelb^-/-^ mutants were detected in the fins of pmela^-/-^; pmelb^-/-^ mutants at 90 dpf. I: In 60 dpf wild-type, pmela^-/-^, pmelb^-/-^ and pmela^-/-^;pmelb^-/-^ caudal fins, the pigmented melanophores in pmela^-/-^ and pmela^-/-^;pmelb^-/-^ were significantly lower than in pmelb^-/-^ mutants and wild-type fish. While the pigmented melanophores in pmelb^-/-^ mutants were also significantly lower than the wild-type fish. In 90 dpf wild-type, pmela^-/-^, pmelb^-/-^ and pmela^-/-^;pmelb^-/-^ caudal fins, the pigmented melanophores in pmela^-/-^;pmelb^-/-^ mutants were significantly lower than in pmela^-/-^ mutants, the pigmented melanophores in pmela^-/-^ mutants were significantly lower than in pmelb^-/-^ mutants, and the pigmented melanophores in pmelb^-/-^ mutants were significantly lower than in wild-type fish. J: The pigmented melanophores in 60 dpf wild-type, pmela^-/-^, pmelb^-/-^ and pmela^-/-^; pmelb^-/-^ mutants were significantly lower than the wild-type fish and mutants at 90 dpf stage, respectively. Data shown in Figure 6I and 6J are expressed as the mean ± SD (n=9). Significant differences in the data between groups were tested by one-way ANOVA and Duncan’s post-hoc test and Student’s t-test, respectively. P<0.05 was considered to be Statistically significant, indicated by different letters above the error bar.

### Body color of *pmel* mutants at different developmental stages

We characterized the phenotype of the mutant fish in their early developmental stages. At 12 dpf, wild-type fish showed no bars, and melanophores were spread evenly across the whole trunk (Fig 7A). At 60 dpf, wild-type fish showed black bars separated by light colored inter-bars. A higher density of melanophores was detected in the bar regions. Iridophore number sharply increased in the inter-bar regions (Fig 7B). At 90 dpf, wild-type fish showed strong black bars separated by iridescent inter-bars (Fig 7C and 7C’), much like the 150 dpf and adult fish (Fig 7M).

**Fig 7.**
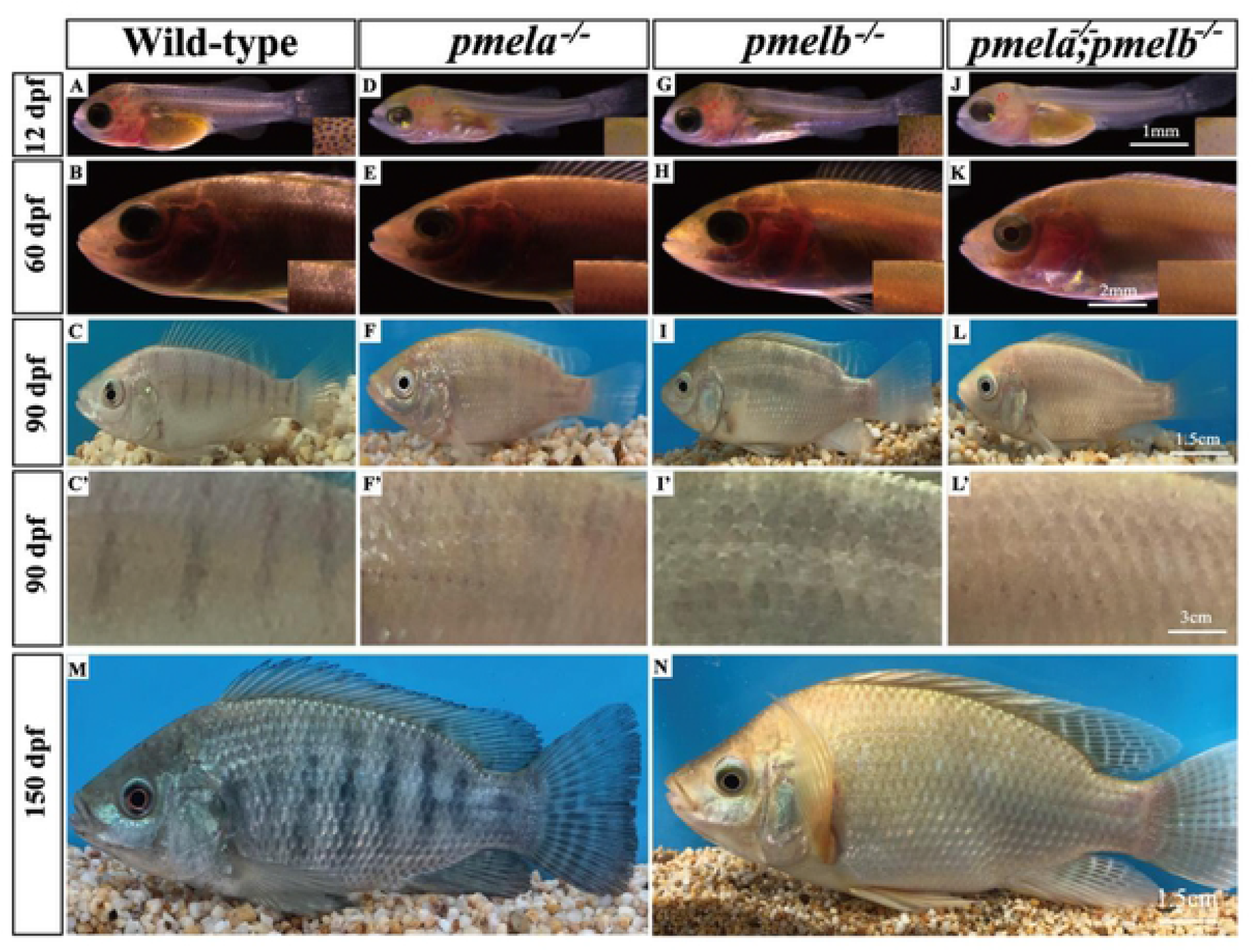
Mutation of pmela and pmelb resulted in serious hypo-pigmentation. The fish were checked at 12, 60, 90 and 150 dpf. Hypo-pigmentation of pmela^-/-^, pmelb^-/-^ and pmela^-/-^; pmelb^-/-^ mutants were observed in trunk at all the developmental stages. A: No bars were observed (the bars usually begin to appear at around 35 dpf) in wild-type fish. B: Black wild-type fish with heavily pigmented vertical bars detected on the trunk at 60 dpf. C and C’:Wild-type fish with obvious vertical bars detected on the trunk at 90 dpf. D: pmela^-/-^ mutants with significant reduction of pigmented melanophores detected in whole fish at 12 dpf. Melanophores in the head and RPE were detected with significant hypo-pigmentation. E: pmela^-/-^ mutants with serious hyper-pigmentation detected in trunk at 60 dpf. No obvious vertical bars were detected at this stage, the body color was evenly detected in the whole fish. Many grey iridophores were detected on the trunk. F and F’: pmela^-/-^mutants with hypo-pigmentation detected in whole fish, which made the fish show light golden body color at 90 dpf. G: pmelb^-/-^ mutants with hypo-pigmentation on trunk, while not on melanophores at 12dpf. H: pmelb^-/-^ mutants with some pigmented melanophores detected in trunk and fins, but many of the melanophores were in abnormal states at 60 dpf. I and I’: pmelb^-/-^ mutants detected with some black patterns which were different with the bars of wild-type fish at 90 dpf. The whole fish showed hypo-pigmentation. J: pmela^-/-^; pmelb^-/-^ mutants detected with even more significant hypo-pigmentation than pmela^-/-^ and pmelb^-/-^ mutants at 12 dpf. The melanophores in the head and RPE were similar to the situation revealed in pmela^-/-^ mutants. K and K’: pmela^-/-^; pmelb^-/-^ mutants were detected with global golden body color, but light grey pigmented melanophores were detected in some areas like the top head at 60 dpf. L and L’: pmela^-/-^;pmelb^-/-^ mutants were detected with golden plumage body color, no bars, but similar light color pattern with pmela^-/-^ and pmelb^-/-^ mutants in the dorsalfin at 90 dpf. M: Wild-type fish with global black plumage and hyper-pigmented vertical bars detected on the trunk and fins at 150 dpf. N: pmela^-/-^;pmelb^-/-^ mutants were detected with complete golden plumage at 150 dpf.

At 12 dpf, the *pmela*^-/-^ mutant fish showed serious hypo-pigmentation across the whole fish (including RPE). Melanophore numbers were reduced on the trunk surface, and the remaining melanophores were light grey, indicative of a reduction in melanin synthesis (Fig 7D). Almost no pigmented melanophores were observed in the mutants at 60 dpf (Fig 7E). Hypo-pigmented patterns on the dorsal fin and a red-black RPE were observed in the lateral view (Fig S10). At 90 dpf, the body color was yellowish, and the iris showed obvious hypo-pigmentation (Fig 7F and 7F’).

At 12 dpf, *pmelb*^-/-^ mutant fish showed hypo-pigmentation but with more pigmented melanophores than the *pmela*^-/-^ mutant fish (Fig 7G). At 60 dpf, melanophores were present in reduced numbers on the trunk surface, and although regular bars were not seen, we occasionally detected irregular patches of pigmentation (Fig 7H). At 90 dpf, the fish showed yellowish body color with slightly pigmented dorsal patches compared with the *pmela*^-/-^ mutant fish (Fig 7I and 7I’).

At 12 dpf, *pmela*^-/-^*;pmelb*^-/-^ double mutants showed a phenotype similar to the *pmela*^-/-^ mutants. The RPE and the trunk were hypo-pigmented. Macro-melanophores with reduced melanin content were observed, but no pigmented melanophores were observed on the trunk (Fig 7J). At 60 dpf, no bars or pigmented melanophores were detected in the double mutants. They showed even more significant hypo-pigmentation compared with the *pmela*^-/-^ or *pmelb*^-/-^ mutants, which gave the whole fish a golden color (Fig 7K). At 90 dpf, significant hypo-pigmentation was detected in the double mutants. No bars or black/grey patterns were detected in the trunk and all the fish showing completely golden color. The iris was hypo-pigmented, but it was difficult to distinguish the differences of RPE pigmentation between the mutants and the wild-type fish from the lateral view (Fig 7L and 7L’). At 150 dpf, the double mutants showed a golden color with reddish fins, and the yellow pigmentation and remaining melanophores were spread evenly across the whole fish. The RPE was as black as the wild-type fish (Fig 7N).

### Pigment cell number, morphology and location were greatly changed in *pmel* mutants

The thin and partially transparent scales as skin appendages provided us excellent materials to investigate the cell number, morphology and location between each type of pigment cells. The 60 dpf (before melanin biosynthesis partially restored) and 150 dpf (melanin gradually accumulate over many weeks) scales of wild-type fish and *pmela*^-/-^*;pmelb*^- /-^ double mutants were used as materials to reveal the relative pigment cell number, morphology and location between the wild-type fish and PMEL-free golden fish. Just as predicted, the number and sizes of melanophores and xanthophores were consistent with the results we showed above in different developmental stages. In wild-type fish, many melanophores were healthy and heavily pigmented with mature melanin, the dendrites were active and fully spread, which made a large branched cell, especially in the older 150 dpf melanophores. Additionally, iridophores were spread mainly on the surface of the mainly body of melanophores. However, the xanthophores were detected with a smaller, round shape. They located near the dendrites of melanophores, or even further away, probably restricted and rejected by melanophores (Fig 8A and 8C). As a contrast, in scales of 60 dpf PMEL-free hypo-pigmented fish, the melanophores with insufficient melanin biosynthesis were often detected with melanin-free melanophores filled with iridescent purple, blue or even red guanine, in both the main body and the dendrites of melanophores (Fig 8B). In the scales of older 150 dpf PMEL-free golden fish, the small number of melanophores detected had a tiny-spot appearance, and the gradually accumulated melanin gathered as a weak spots, with a much smaller size than that of the wild-type melanophores. Additionally, the xanthophores were obviously enlarged with “airenemes” in the mutants compared with those of the wild-type. The larger sizes of xanthophores allowed the mutants to produce more yellow pteridines/carotenoids. Besides, the xanthophores located near the melanophores, and some of them even clung on the main body of the tiny-spot-like melanophores, indicating that the nearest-neighboring distances between the melanophores and xanthophore were probably reduced and the interactions of them were heavily influenced. The remaining iridophores were arranged in a line between the weak melanophores, and eventually combined into a fish net structure in the scales (Fig. 8D). These results suggeste that the cell-cell interactions between the pigment cells (both the same type and different types of pigment cells) were altered in the PMEL-free golden fish. The relative pigment cell number, morphology and location (mainly between the melanophores and xanthophores) were completely different between the wild-type and mutants, which was the fundamental reason for the formation of golden body color in tilapia.

**Fig 8.**
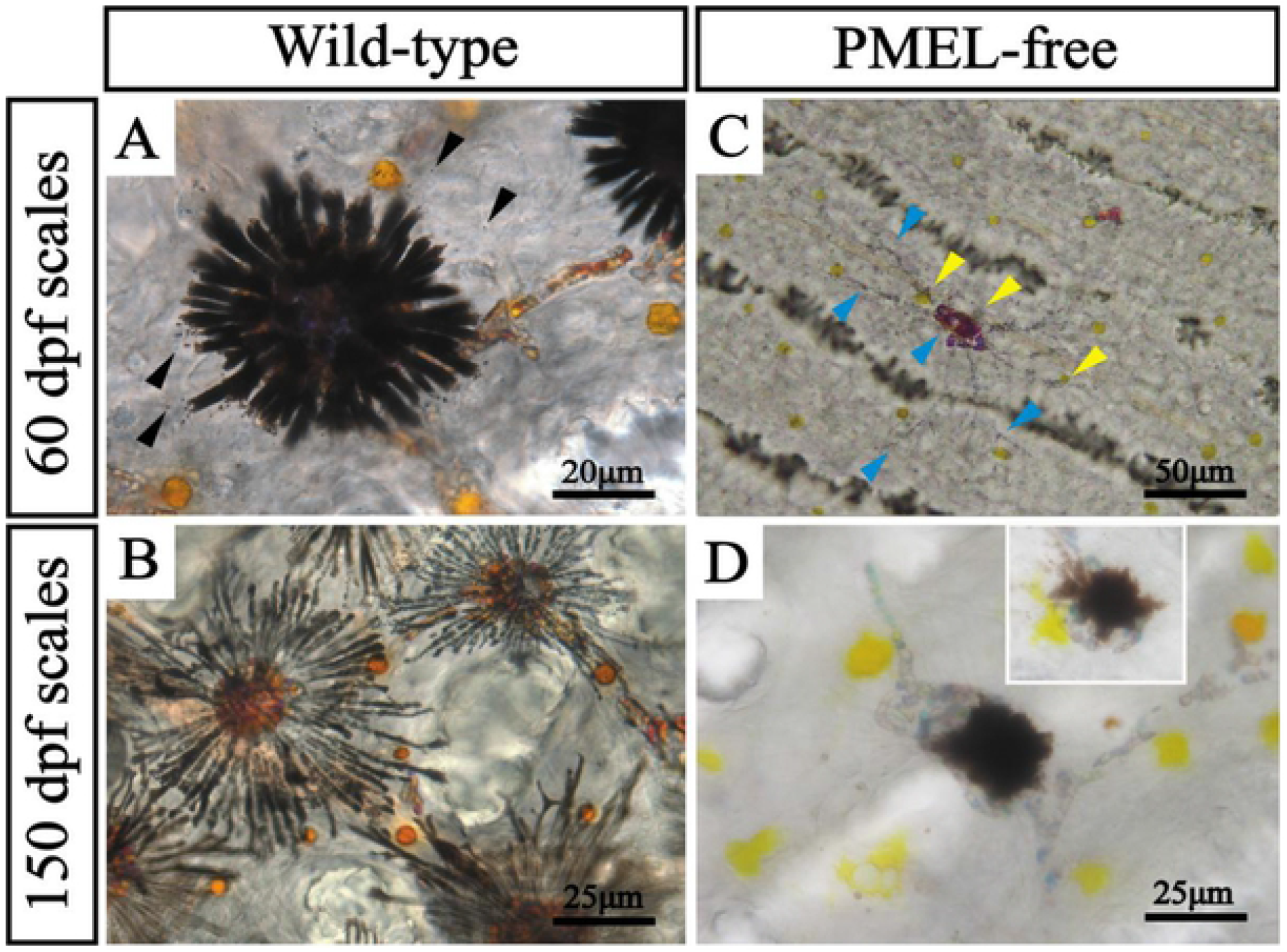
Effects of pmela and pmelb double mutation on pigment cell number, morphology and location in scales of tilapia. In the wild type, most of the melanophores were young, with many dendrites and extracellular release of melanosomes at 60 dpf (A, black arrow). With the development, most of the melanophores became elder. The increased and further extended dendrites propelled the xanthophores far away from the main body of the melanophore at 150 dpf (B). In pmela^-/-^;pmelb^-/-^ mutants, the main body and dendrites of the melanophore shrank, and accumulated a large amount of purple, blue and even red guanine due to the decreased ability of melanin synthesis. The xanthophores became closer to the melanophores at 60 dpf (C, yellow arrow). At 150 dpf, the main body, but not the dendrites, was filled with melanosome, due to the partially recovered melanin synthesis ability. Xanthophores were obviously enlarged with “airememes” in the mutants compared with those of the wild-type (D).

## Discussion

### *Pmela* and *pmelb* showed similar expression patterns in Nile tilapia

*Pmel* is a key gene involved in melanin biosynthesis and the development of body color in many fish species, such as carp [17, 36], and cichlids [24, 37, 38]. In those studies, the expression levels of *pmela* in hyper-pigmented skin were always significantly higher than in hypo-pigmented skin. In the skin transcriptome of Malaysian red tilapia, *pmela* was found to be expressed significant higher in black skin than in red/pink skin [37], suggesting that *pmel* might be involved in skin color differentiation between black and red tilapia. Additionally, this study suggested that *pmela* and *pmelb* were most highly expressed in eyes, indicating that the two genes might be involved in eye pigmentation and development. In zebrafish, *pmela* has been shown to be involved in eye pigmentation and anterior segment size maintenance in early juvenile stages, through loss of function studies [18]. In different developmental stages of wild-type Nile tilapia, *pmela* and *pmelb* were detected with similar expression patterns. They were both highly expressed in early developmental stages (from 6-12 dpf), which was in consistent with the several waves of melanophore proliferation identified in our previous studies of color patterning in Nile tilapia [33]. However, no expression of *pmela* and *pmelb* was detected at 2-4 dpf, probably because these genes are downstream in the melanogenesis pathway, and follow the expression of NCCs-melanophores specification genes like *mitf* and *kita/kitlga*. Moderate expression of *pmela* and *pmelb* was also detected in fish at 30-90 dpf and the adult stage, suggesting that they are probably involved in melanin biosynthesis or melanophore survival at later stages. The *pmelb* was detected with some expression in dorsal fins in adult wild-type fish, which was in accordance with the results of our loss of function studies. The *pmela*^-/-^ mutants were still observed with banding in dorsal fin at 60 dpf, while *pmelb*^-/-^ mutants presented a more serious hypo-pigmentation of this fin. This was similar to the expression analysis in the cichlid, *Neolamprologus meeli*, in which higher *pmel* expression was detected in adult dorsal fin than in the ventral part of the anal fin or the caudal fin [39].

### Homozygous mutation of *pmel* genes resulted in golden color in tilapia

Using CRISPR/Cas9 gene editing, we successfully disrupted the expression of PMEL. Different levels of hypo-pigmentation were detected in the F0 mutants [33]. As predicted, the homozygous mutants of *pmela, pmelb* and *pmela;pmelb* all showed obvious hypo-pigmentation. All three mutants had yellowish/golden body color, especially in the double mutants. The *pmela*^-/-^*;pmelb*^-/-^ double mutants had completely golden bodies, without any black/grey patches or spots, which made them quite attractive (Fig 7N and Fig S9). However, the golden fish had pigment patterning in fins. This was similar to studies in zebrafish, in which disruption of *csf1ra*, a xanthophore differentiation marker, led to different phenotypes in trunk and fins [40]. Additionally, studies of pigment cells in *Danio* and other teleosts also suggested that color patterns in trunk and fins are probably controlled by different genes [26, 41]. In a recently published manuscript, the gene *pmel17* was found to be closely linked with color traits in a naturally mutant yellow Mozambique tilapia population [42].

### The effects of PMEL mutation on pigment cells relative abundance and melanin biosynthesis in Nile tilapia

As mentioned above, loss of function of both *pmela* and *pmelb* in Nile tilapia led to a complete golden skin color without any black bars or patches, but with hypo-pigmented eyes, due to the increased numbers and sizes of xanthophores, and decreased numbers and sizes of melanophores, as well as hypo-pigmentation reflected by deficient in melanin biosynthesis (Fig 8, 9A and 9B).

**Fig 9.**
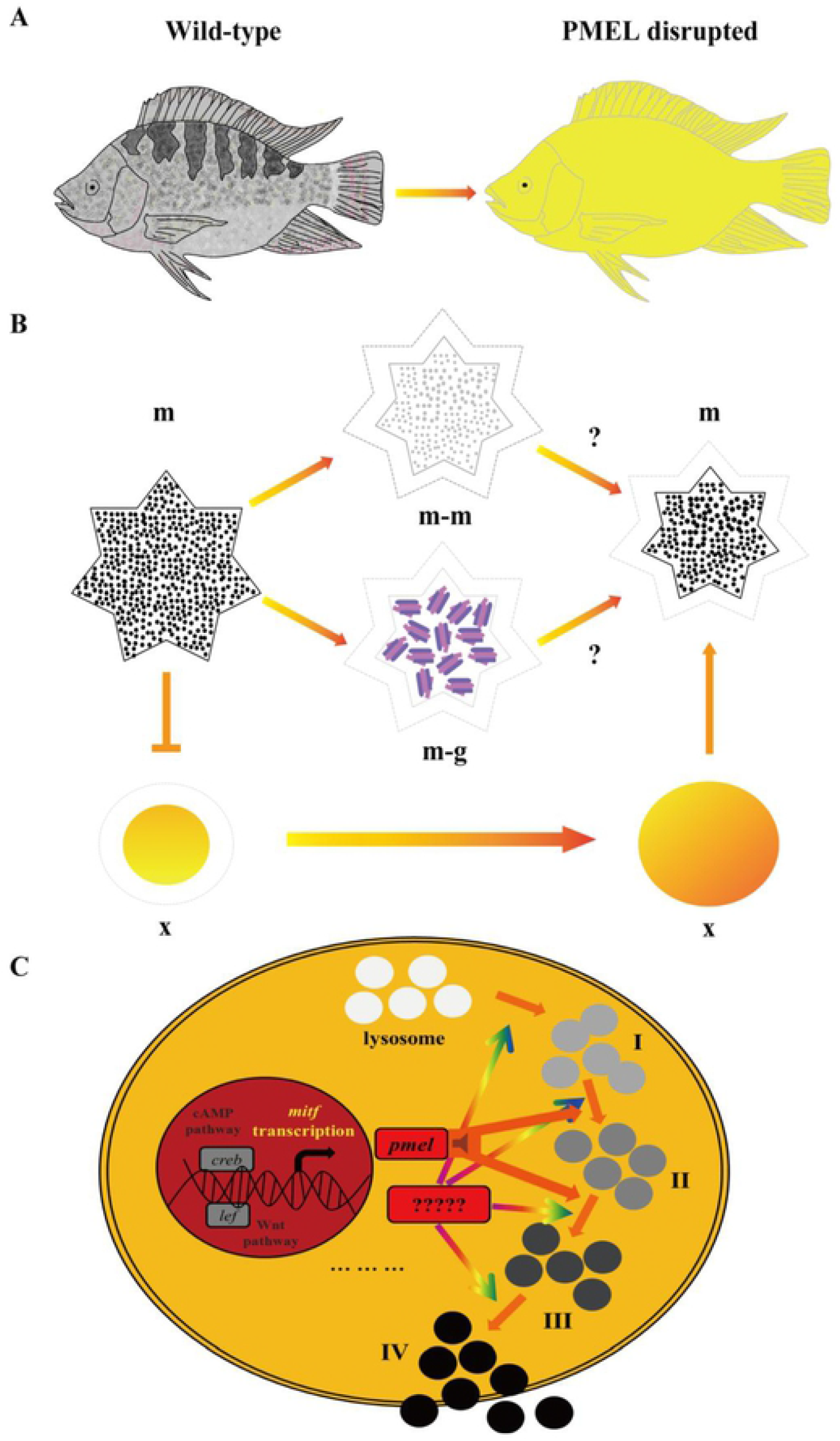
Proposed model of PMEL disruption in this study. A: Disruption of pmela and pmelb resulted in complete golden plumage and hypo-pigmentation in eyes of tilapia. B: Disruption of pmela and pmelb resulted in decreased and hypo-pigmented melanophores, and even guanine biosynthesis in not pigmented melanophores. Additionally, melanophores size was also heavily reduced in it. Disruption of pmela and pmelb also resulted in increased and larger sized xanthophores, which was another key reason for complete golden plumage. In wild-type fish, larger sized melanophores rejected the size-restricted xanthophores; while in PMEL-free mutants, the xanthophores with larger size than in wild-type fish pursuited the functionally restricted tiny melanophores. C: Schematic view of melanin biosynthesis. Mitf as the key transcription factor for melanogenesis is responsible for cis-directing the mitf-axis down stream genes such as pmel. There are four phases of melanin, pmel was fundamental for both phase I to phase II and phase II to phase III melanin transition, thus hypo-pigmented melanophores were observed in PMEL disrupted tilapia. However, restoration of melanin biosynthesis were observed in PMEL disrupted tilapia, suggesting that additional collateral branches and some other key genes might be involved in melanin biosynthesis. The unknown gene was probably related to melanin biosynthesis from phase I to phase IV, and even lysosome biosynthesis. m, melanophores; m-m, melanophores with immature melanin; m-g, melanophores fulfilled with guanine; x, xanthophores.

Some of the melanin-free melanophores were able to accumulate guanine, especially in early developmental stages (stages before 60 dpf). During the time period the pigmentary basis for the continued presence of banding in the fins of mutant fish were bands of guanine. We did not detect red erythrophores until at least 50 dpf in our previous studies [33], and the sizes and shapes were most similar to melanophores. The 60 dpf wild-type fish also showed many melanophores with different levels of black/grey melanin. Previous studies on zebrafish suggested that melanophores were still detected in *tyr* homozygous mutants, but they did not contain melanin pigment [43]. Our results are similar to those from zebrafish *tyr* mutants, but the tilapia PMEL-free mutants displayed milder defects compared to the zebrafish *tyr* mutants.

Melanophores size was also affected by mutation of *pmel*, as melanophores were smaller in the mutants, not only in the whole trunk and fins, but also in the scales (Fig 8D and 9B). This is likely to be a result of defects in forming a fibrillar structure within the melanosome upon which melanin is deposited [44, 45], which finally led to the appearance of small melanophores with insufficient melanosome development.

At the same time, mutation of *pmel* genes caused increases in the numbers and sizes of xanthophores, which plays a decisive role in the complete golden phenotype. In wild-type fish, xanthophores are small and round in shape, while the neighboring melanophores are dendritic. In *pmela*^-/-^*;pmelb*^-/-^ mutants, xanthophores were larger than in wild-type fish, while melanophores were detected with spot-like shapes, not only in the trunk surface and fins, but also in the scales (Fig 8C, 8D and 9B).

*Mitf* is a key transcription factor for melanin biosynthesis, and is necessary for all the processes from the development of melanosomes to mature melanin release. Both *pmel* genes have promoter sequences that suggest they are regulated by *mitf*. Probably *pmel* was critical for melanosome transition from phase I to III, as we detected hypo-pigmented melanophores with light grey melanin in PMEL-disrupted fish. However, we detected slow restoration of melanin biosynthesis during development (Fig 9C), possibly because some melanin synthesis can occur in the absence of the fibrillar substrate provided by PMEL. Even though melanin could accumulate gradually over many weeks, the mutants did not produce a melanin-pigmented phenotype as the melanophores were small.

### *Pmela* was more important than *pmelb* in RPE pigmentation and eye development

In this study, both *pmela* and *pmelb* were detected with the highest expression levels in the eyes of wild-type fish. They were further proved to be involved in eye pigmentation by our loss of function studies. Significant less global pigment was detected in the eyes of *pmela*^-/-^ and *pmela*^-/-^*;pmelb*^-/-^ mutants. *Pmela* was further confirmed to be necessary for eye pigmentation (hypo-pigmentation of the RPE and iris) in early stages of eye development, which was in consistent with the results from human and zebrafish [18]. In a phylogenetic analysis across different fish species, *pmelb* was suggested to be responsible for adaptive phenotypes (specifically melanin-free in both eyes and the whole trunk, and eye degeneration) in extreme environments [12]. In our studies, disruption of *pmela* resulted in RPE pigment loss. The *pmela*^-/-^ and *pmela*^-/-^*;pmelb*^-/-^ mutants had a deep red RPE and also a hypo-pigmented iris, while the *pmelb*^-/-^ mutants had an almost normal pigmented RPE, but a hypo-pigmented iris. The results indicate that both *pmela* and *pmelb* are critical for iris pigmentation, while *pmela* played a more important role than *pmelb* in RPE pigmentation and eye development in tilapia. Whether this is true in other teleosts species remains to be investigated.

### Both *pmela* and *pmelb* were important for body color formation in Nile tilapia

Although many studies of gene expression suggested that *pmel* might function in body color differentiation [17, 24, 36-38], ours is the first detailed functional analysis of *pmela* and *pmelb* in teleosts. The effects of *pmel* mutations appear to vary among species [19, 20]. *Pmel* mutation has been confirmed to be the main cause of eye abnormal development including RPE pigment loss and anterior segment abnormal development in human beings and zebrafish [18]. The phenotypes of *pmela* mutants in tilapia, hypo-pigmentation of trunk surface and RPE, were similar to those revealed in larvae zebrafish.

Body color is an important economic trait. In aquaculture, the fish with pleasant body color is always preferred by consumers. For example, the koi carp and golden fish have been raised as pet fish all over the world, due to their attractive body color or amazing specific color patterns. In tilapia, the red tilapia is also preferred by consumers globally because of the lack of black pigmentation in the trunk and peritoneum [46, 47]. The establishment of a golden tilapia (*pmela*^-/-^*;pmelb*^-/-^ mutants) would be of great significance to the tilapia aquaculture industry, and might also be of interest in the pet fish market.

## Materials and methods

### Fish

The founder strain of Nile tilapia was obtained from Prof. Nagahama (Laboratory of Reproductive Biology, National Institute for Basic Biology, Okazaki, Japan). This strain has been in laboratory culture for more than 20 years, and is thus largely homozygous. The founder strain of red tilapia in our lab is Malaysia red tilapia which was obtained from Hainan fish farm. Experimental tilapia were reared in recirculating aerated freshwater tanks and maintained at ambient temperature (27 °C) under a natural photoperiod. Prior to the experiments, the fish were kept in laboratory aquariums under 15:9 h light: dark conditions at temperature of 27±1 °C for one week. All animal experiments conformed to the Guide for the Care and Use of Laboratory Animals and were approved by the Committee for Laboratory Animal Experimentation at Southwest University, China.

### Identification of *pmel* genes from different animal species

We examined the genomes of 15 animal species (zebra cichlid *Maylandia zebra*, Nile tilapia *Oreochromis niloticus*, Japanese medaka *Oryzias latipes*, guppy *Poecilia reticulata*, torafugu *Takifugu rubripes*, channel cat fish *Ictalurus punctatus*, spotted gar *Lepisosteus oculatus*, zebrafish *Danio rerio*, tongue sole *Cynoglossus semilaevis*, coelacanth *Latimeria chalumnae*, common frog *Rana temporaria*, human *Homo sapiens*, house mouse *Mus musculus*, chicken *Gallus gallus* and common lizard *Zootoca vivipara*) to identify *pmel* genes in each species. These 15 species were representative animal species from fishes and also groups at different evolutionary position. The genomic sequences of all species are available at the NCBI and Ensemble data base. They have relatively high quality genome sequences which allow us to isolate both *pmel* genes to reflect the true evolutionary history. All *pmel* genes were identified by tblastn (E=2×10−5) against genome sequences, using zebrafish PMELA and PMELB proteins.

### Phylogenetic analyses and genomic distribution

We chose to use more conservative amino acids sequences, rather than nucleotide sequences for phylogenetic analyses. The amino acid sequences of *pmel* genes were aligned by ClustalW with default parameters using the multiple alignments of software BioEdit [48]. Phylogenetic trees were generated by the neighbor joining (NJ) method using the program MEGA6.0 software [49]. The final picture of NJ tree was modified with Adobe Illustrator CS6 (Adobe Inc. USA).

### The gene functional domain prediction and promoter binding sites analysis

The functional domain prediction of tilapia PMELA and PMELB was carried out in SMART online program. The 2k bp promoter sequences of *pmela* and *pmelb* were downloaded from the NCBI database. The cis-regulating elements of the promoter were predicted and analyzed in AnimalTFDB3.0 program, and the MITF binding consensus sequences were highlighted.

### RT-PCR validation of *pmela* and *pmelb* expression

The *pmela* and *pmelb* specific primers used for reverse transcription PCR amplification were designed using Primer Premier 6. The sequences are listed in Supplemental Table 1. Triplicate samples were collected at 2, 4, 6, 8, 10 dpf (whole fish) and 20, 30, 60, 90 dpf (skin) and adult stage (different tissues, including skin). Reverse transcription was conducted in a total reaction volume of 20 μl, which included 2 μg total RNA and 2μl RT reaction mixture. For PCR amplification, *pmel* specific primers or *β*-actin primers were added into the reaction at the beginning of PCR and each PCR run for 34 cycles. The PCR products were separated by agarose gel electrophoresis and photographed under UV illumination.

### Establishment of *pmela*^-/-^, *pmelb*^-/-^ and *pmela*^-/-^*;pmelb*^-/-^ mutants by CRISPR/Cas9

The *pmela* and *pmelb* mutant fish with the highest indel frequency were used as G0 founders. They were raised to sexual maturity and mated with wild-type tilapia. F1 larvae were collected at 10 dah and genotyped by PCR amplification and subsequent *Fsp*EI and *Tsp*45I digestion.

CRISPR/Cas9 was performed to knockout *pmela* and *pmelb* in tilapia as described previously [33]. Briefly, the guide RNA and Cas9 mRNA were co-injected into one-cell-stage embryos at a concentration of 150 and 500 ng/µL, respectively. Twenty injected embryos were collected 72 h after injection. Genomic DNA was extracted from pooled control and injected embryos and used to access the mutations. DNA fragments spanning the target site was amplified. The mutated sequences were analyzed by restriction enzyme digestion with *Fsp*EI and *Tsp*45I and Sanger sequencing.

Heterozygous F1 offspring were obtained by F0 XY male founders mated with WT XX females. The F1 fish were genotyped by fin clip assay and the individuals with frame-shift mutations were selected. XY male and XX female siblings of F1 generation, carrying the same mutation, were mated to generate homozygous F2 mutants. The *pmela*^-/-^, *pmelb*^-/-^ and *pmela*^-/-^*;pmelb*^-/-^ mutants were screened using restriction enzyme digestion and Sanger sequencing. The genetic sex of each fish was determined by genotyping using sex-linked marker (marker 5) as described previously [50].

### Image recording and pigment cell observation at different developmental stages

Larvae fish at 5, 7, 12 and 30 dpf and early juvenile stage at 60 dpf were shifted to an observation dish with clean water, photographed by Olympus SZX16 stereomicroscope (Olympus, Japan) under bright or transparent field with different magnification. The 90 dpf wild-type and mutant fish were shifted to the same 15 × 5 × 15 cm^3^ glass water tanks separately, before being photographed with a Nikon D7000 digital camera (Nikon, Japan) against a blue background. ACDSee Official Edition software (ACDSystems, Canada) and Adobe Illustrator CS6 (Adobe Inc. USA) were used to format the pictures.

### Pigment cell analysis of the *pmela*^-/-^, *pmelb*^-/-^ and *pmela*^-/-^*;pmelb*^-/-^ mutants

Larvae submerged in clean water at 7 and 12 dpf were photographed from the lateral view by Olympus SZX16 stereomicroscope (Olympus, Japan) under bright field. Caudal fin from fish at 60 and 90 dpf were photographed with a Nikon D7000 digital camera (Nikon, Japan) against a white background. The caudal fins were removed with medical scissors, soaked in 0.65% Ringers’ solution and directly observed with Olympus SZX16 stereomicroscope (Olympus, Japan) without cover slip under transparent or bright field. Scales from fish at 60 dpf and 90 dpf were soaked in 0.65% Ringers’ solution under cover slip, and were observed under microscope (Germany, Leica EM UC7). Image recording of pigment cells was conducted as quickly as possible after putting them in the Ringers’ solution as the preparations are not stable. ACDSee Official Edition software and Adobe Illustrator CS6 were used to format the pictures. To analyze the number of melanophores, nine fish per group were anesthetized with tricaine methasulfonate (MS-222, Sigma-Aldrich, USA) and immersed in 10 mg/ml epinephrine (Sigma, USA) solution for 15 min to contract melanin. The sizes of the pigment cells and global pigmentation were measured using Image J software [51]. GraphPad Prism 5.01 software (Graphpad, USA) was used to analyze the differences in the numbers and sizes of melanophores in *pmela*^-/-^, *pmelb*^-/-^ and *pmela*^-/-^*;pmelb*^-/-^ mutants and wild type fish. Data values (mean ± SD) were statistically evaluated by one-way ANOVA with Duncan’s post-hoc test and Student’s *t*-test. P<0.05 was considered to be statistically significant, as indicated by different letters above the error bar.

## Supporting information

**Fig S1. There are six putative MITF binding sites in the promoter region of *pmela*, predicted by AnimalTFDB 3**.**0 at**

**http://bioinfo.life.hust.edu.cn/AnimalTFDB/#!/tfbs_predict**.

(TIF)

**Fig S2. There are five putative MITF binding sites in the promoter region of *pmelb*, predicted by AnimalTFDB 3**.**0 at**

**http://bioinfo.life.hust.edu.cn/AnimalTFDB/#!/tfbs_predict**.

(TIF)

**Fig S3. Tissue distribution of *pmela* and *pmelb* in Nile tilapia**. Both *pmela* and *pmelb* expressed highly in eyes, skin and whole embryo. The whole expression level of *pmela* was higher than *pmelb*. Data downloaded from NCBI database [52].

(TIF)

**Fig S4. Tissue distribution of *pmela* and *pmelb* in Nile tilapia**. Both *pmela* and *pmelb* were highly expressed in eyes, and *pmela* was expressed in testis, but not in other tissues examined. At the same time, *pmelb* not only expressed in eyes, but also expressed in the brain and fins, while not in other tissues. B: brain. P: pituitary. HK: head kidney. H: heartoûSP: spleen. I: intestine. K: kidney. L: liver. T: testis. O: ovary. E: eyes. G: gill. M: muscle. SK: skin. F: fins.

(TIF)

**Fig S5. Ontogeny expression of *pmela* and *pmelb* in the skin of different developmental stages in tilapia analyzed by RT-PCR**. *Pmela* was detected with obvious expressions at every developmental stages except for 2 dpf stage. The highest expressions of *pmela* were detected at 8 dpf and 10 dpf stages. A milder expression of *pmela* was detected at 90 dpf. Low expressions of *pmela* were detected with at 4, 6, 20, 30, 60 dpf and adult stages. No expressions of *pmelb* was detected at 2, 30, 60 dpf and adult stage, but high expressions were detected at 4, 6, 8 and 10 dpf. Low expressions of *pmelb* were detected with at 20 and 90 dpf stages.

(TIF)

**Fig S6. Establishment of *pmela***^-/-^***;pmelb***^-/-^ **mutant lines**. *Pmela;pmelb* F2 generation were obtained by firstly mating male F1 *pmela*-positive fish (−5 bp) with female F1 *pmelb*-positive (−7 bp) fish, then heterozygous *pmela;pmelb* F1 offspring with a -5 bp deletion in *pmela* and -7 bp deletion in *pmelb* were selected to breed the F2 generation. The *pmela;pmelb* homozygous mutants accounts for 1/16 in the total F2 offspring.

(TIF)

**Fig S7. Melanophore releases melanosome extracellularly in scales of tilapia**. Like the tetrapods, melanophores in teleosts were also able to release the melanosome extracellularly. The black arrows show the melanosome released.

(TIF)

**Fig S8. Non-pigmented melanophores biosynthesize/accommodate guanine in scales of *pmela***^-/-^***;pmelb***^-/-^ **mutant tilapia at 60 dpf**. The figure showed the scales of PMEL-free mutants at 60 dpf, in which a single melanophore was fullfied with iridescent purple, blue and even red guanine, both in the mainbody and branching-like clusters.

(TIF)

**Fig S9. Mutation of *pmela;pmelb* resulted in hypo-pigmented skin at 90 dpf**. The *pmela*^-/-^ *;pmelb*^-/-^ mutants were yellowish with very serious hypo-pigmentation across the whole fish. However, no significant differences of the RPE pigmentation were detected by the naked eyes.

(TIF)

**Fig S10. Mutation of *pmela* resulted in serious hypo-pigmentation in tilapia at 60 dpf**. The wild-type fish was grayish black with obvious black vertical bars. No bars and pigment patterns were detected in the trunk, pectoral, pelvic, anal and caudal fins of the homozygous mutant. Pigment patterns similar to the wild type fish were detected in the dorsal fin of the mutants. The iris of mutants was silvery white, melanin synthesis was obviously insufficient, and the RPE was dark red.

(TIF)

**S1 Table. Sequences of primers used in the present study**.

(XLSX)

## Author Contributions

**Conceptualization:** Chenxu Wang, Deshou Wang, Thomas D. Kocher.

**Data curation:** Chenxu Wang, Deshou Wang, Thomas D. Kocher, Jia Xu, Minghui Li.

**Formal analysis:** Chenxu Wang, Jia Xu, Thomas D. Kocher, Deshou Wang.

**Funding acquisition:** Deshou Wang, Chenxu Wang, Minghui Li, Thomas D. Kocher

**Investigation:** Chenxu Wang, Minghui Li, Thomas D. Kocher.

**Methodology:** Chenxu Wang, Feilong Wang, Zhuo Yang, Minghui Li, Jia Xu.

**Project administration:** Deshou Wang, Chenxu Wang, Thomas D. Kocher, Minghui Li.

**Resources:** Deshou Wang, Chenxu Wang.

**Software:** Chenxu Wang, Feilong Wang.

**Supervision:**

**Validation:** Chenxu Wang, Deshou Wang, Thomas D. Kocher,.

**Visualization:** Chenxu Wang, Thomas D. Kocher, Deshou Wang.

**Writing – original draft:** Chenxu Wang, Deshou Wang, Thomas D. Kocher.

**Writing – review & editing:** Chenxu Wang, Deshou Wang, Thomas D. Kocher, Jia Xu, Minghui Li.

## Data Availability Statement

The required links or identifiers for our data are present in the manuscript as described.

